# Systematic data-driven genome-scale metabolic model reduction for dynamic bioprocess modeling: CHO cell culture case study

**DOI:** 10.1101/2025.10.02.679947

**Authors:** Athanasios Antonakoudis, Anne Richelle

## Abstract

Genome-scale metabolic models (GEMs) enable mechanistic insight into cellular metabolism, but their size and underdetermination hinder use in dynamic bioprocess simulation and real-time digital twins. Compact models are essential, yet existing reduction strategies either neglect experimental uncertainty, rely on simplistic rate estimates, or depend on manual assumptions, limiting robustness and scalability. Here, we present a metabolomics-driven reduction pipeline that integrates Bayesian flux estimation to propagate uncertainty from noisy and sparse exo-metabolomics data directly into the reduction process. Applied to time-course data from 12 fed-batch CHO cultures, the method produced a single reduced model that remained feasible across all conditions, avoided over- and under-pruning, and accurately reproduced observed extracellular fluxes. Despite relying solely on exo-metabolomics, the reduced model preserved broad metabolic functionality, highlighting the strong predictive power of extracellular data. This establishes a systematic, uncertainty-aware framework for generating compact GEMs suited for dynamic bioprocess simulation and digital twin integration, demonstrated here in a CHO case study but generalizable across cell systems.

## 1. Introduction

Genome-scale metabolic models (GEMs) provide comprehensive representations of cellular metabolism, enabling integration of omics data and prediction of metabolic phenotypes through flux balance analysis (FBA) and related methods (Oberhardt et al., 2009). They have become central to strain design in microbes, the study of human metabolism and disease, and the integration of multi-omics data, with applications extending to mammalian cell systems and bioprocesses (Gu et al., 2019; Strain et al., 2023). However, despite this broad utility, GEMs are rarely the primary choice for bioprocess modelling or digital twin development due to their large size and degrees of freedom, which make real-time use impractical (Richelle et al., 2020; Park et al., 2024; Ranpura et al., 2025). As a result, digital twins are more often constructed from empirical or statistical models, black-box machine learning, or deterministic mechanistic expressions based on ordinary differential equations (ODEs). These approaches can be parameterised quickly and executed efficiently for integration with process analytical technologies (PAT) (Chen et al., 2020; Kreuzer et al., 2024).

The challenge with GEMs is that they encode the full metabolic repertoire of an organism, including pathways inactive under the conditions of interest. While this comprehensiveness is advantageous for systems biology, it makes the models inherently large and underdetermined, requiring extensive data to constrain. In bioprocess settings, however, data are typically sparse and noisy, leaving GEMs insufficiently constrained and prone to unstable or infeasible predictions. Several model reduction strategies have been developed to address the size and complexity of GEMs, with metabolomics-guided approaches being among the most widely applied. These methods infer metabolite uptake and secretion rates from experimental concentration profiles and use them to constrain flux distributions, thereby identifying and deactivating reactions unlikely to carry flux under the studied conditions. Despite their utility, these approaches face a fundamental limitation: they typically treat estimated rates as exact values within sampling intervals, neglecting both measurement uncertainty and the dynamic variability of cellular metabolism (Bernstein et al., 2021). As a result, strict use of uncertain rate estimates can cause both over-pruning, by discarding pathways that should have remained feasible, and under-pruning, by retaining reactions that should be inactive. These distortions compromise the predictive reliability of reduced models in dynamic bioprocess simulations.

Alongside omics-guided strategies, algorithmic reduction frameworks such as redGEM (Ataman et al., 2017), NetworkReducer (Erdrich et al., 2015), and SWIFTCORE (Tefagh and Boyd, 2020) adopt a network-structural perspective on model compression. By systematically removing or merging reactions while enforcing flux consistency, these methods can generate compact subnetworks without relying on extensive experimental rate measurements. They often do so by pre-defining a core set of reactions or lumping biomass precursors into simplified pathways, thereby retaining a minimal yet flux-balanced scaffold. While such reductions are well-suited for static simulations or exploratory analyses, their utility in dynamic bioprocess contexts has limitations. The reliance on strong a priori assumptions about pathway essentiality may bias outcomes, and the lumping of reactions can obscure biological interpretability. More importantly, feasibility is typically validated only under fixed conditions, leaving the reduced models ill-equipped to accommodate uncertainty, respond to changing process environments, or assimilate new data streams - all critical requirements for bioprocess digital twins.

Several studies have attempted to adapt GEM reductions for CHO cell bioprocesses, each addressing important aspects but leaving critical gaps unresolved. Martínez et al. (2015) highlighted the importance of accurate rate estimation by using B-spline fitting to capture dynamic flux changes during temperature shifts. However, the reduced model was constructed separately from the data and not directly guided by the calculated rates, limiting its integration with bioprocess simulations. Ramos et al. (2022) developed a medium-specific reduction to support their HybridFBA framework, combining stoichiometric constraints with PCA-derived correlations to improve predictive power. While this represents a semi-systematic reduction approach, it relied on constant rate estimates and imposed an a priori noise, limiting its ability to robustly handle experimental uncertainty. Calmels et al. (2019) demonstrated the industrial applicability of reduced CHO GEMs in fed-batch processes, but the model was manually curated and constrained using traditional constant rate estimates, raising questions of scalability and reproducibility. More recently, Ghodba et al. (2025) integrated kinetic rate laws into a CHO GEM to simulate fed-batch dynamics. While this advanced the treatment of rate estimation, the reduction was minimal - limited to the removal of blocked reactions - leaving the network highly underdetermined and challenging to use for reliable prediction.

Together, these studies underscore both the potential and the remaining limitations of reduced GEMs in bioprocess applications. Some emphasize rate estimation but overlook systematic reduction; others reduce model size but rely on simplistic or noisy rate assumptions; and industrial applications often depend on manual curation, limiting scalability. A unifying limitation is the lack of uncertainty-aware integration of rate estimation with model reduction, which is essential for feasibility and robustness in dynamic digital twin contexts. More broadly, recent perspectives on digital biomanufacturing highlight that full GEMs are computationally intractable for online integration, reinforcing the need for compact, systematically validated models to enable real-time bioprocess control (Monteiro et al., 2023; Ranpura et al., 2025).

To address this need in industry, we present an exo-metabolomics-driven reduction pipeline specifically designed for dynamic bioprocess simulation. The approach integrates Bayesian-derived extracellular flux estimates that provide uncertainty-aware rate bounds (Andersson et al., 2025; Richelle et al., 2025), producing smoother and more stable constraints than conventional constant rate or spline-based methods. These probabilistic bounds are used to systematically guide the reduction of the iCHO1766 model (Hefzi et al., 2016), with explicit feasibility checks at every step to ensure solvability across conditions and time points. This framework directly addresses the limitations of prior work: it couples rate estimation with reduction, propagates experimental uncertainty instead of imposing arbitrary noise assumptions, and avoids over- or under-pruning by operating on probabilistic bounds. Importantly, the resulting reduced model preserved core metabolic functionality and broad gene coverage, comparable to transcriptomics-guided reductions but without requiring additional omics inputs or manual curation. Together, this establishes a robust and systematic pipeline for generating compact GEMs tailored to dynamic bioprocess simulation and real-time digital twin integration.

## 2. Material and Methods

### 2.1 Cell culture data and experimental measurements

Chinese Hamster Ovary (CHO) DG44 cells (Sartorius) producing an IgG1 monoclonal antibody were cultured in chemically defined Sartorius medium. Twelve fed-batch cultures were performed in Ambr® 250 bioreactors (200 mL working volume) for 14 days, inoculated at 0.3 × 106 cells/mL. Fed-batch cultures were performed with a production medium (PM-Sartorius) and with two feed media (FMA and FMB-Sartorius). The feeding profiles were calculated offline following the standard applications implemented in Sartorius, and the corresponding amounts of each feed media were added daily after Day 3. Also, a daily glucose bolus was performed starting on day 5 of culture when the measured glucose concentration was less than 5 g/L (stock glucose solution of 400 g/L). Cultures were maintained at 36.8 °C, pH 7.1, and 60% dissolved oxygen, with agitation adjusted to maintain DO and automated antifoam addition. Temperature or pH shifts were introduced at day 7 to generate distinct process conditions. Each batch was sampled daily, measuring viable cell concentration, viability, product secretion, and concentrations of ∼60 extracellular metabolites using a combination of NMR spectroscopy, enzymatic assays (FLEX), and LC–GC methods.

### 2.2 Rate calculation using Bayesian inference

Metabolic rates were quantified utilizing the MetRaC framework, which integrates Bayesian inference with nested sampling techniques (Andersson et al., 2025; Richelle et al., 2025). This approach first adjusts raw concentration profiles to account for variations in reactor volume, such as those resulting from feeding events and sampling procedures. Subsequently, it models consumption and/or production dynamics using a linear combination of basis functions (i.e., logistic function). Bayesian parameter estimation via nested sampling yields posterior distributions for the model parameters, facilitating uncertainty quantification and providing a robust foundation for model comparison. This approach produces an ensemble of rate trajectories that capture the full posterior distribution of the estimates. Rates can then be sampled at arbitrary time points, effectively increasing the number of usable experimental data points while preserving uncertainty. For this study, 100 trajectories were generated per culture, each sampled at 100 evenly spaced time points across the cultivation period. At each sampling time, the median as well as the 5th and 95th percentiles of the rate distribution were computed and used as inputs for the reduction strategy.

### 2.3 Constant rate calculation

To establish a baseline comparison with the MetRaC-derived rates, we employed a more traditional method, referred to here as the “constant rate approach”, for the calculation of metabolic rates at every time point. The uptake or secretion rate of a metabolite *M* at time *i* was estimated as:

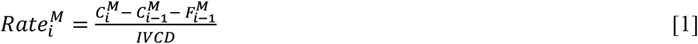

where, *C*_*i,i*−1_ are the metabolite concentrations at times *i* or *i* − 1, *F* is the feeding contribution, and *IVCD* is the integral of viable cell density. The constant rate calculation provides a single-point estimate of the metabolic rate at time *i*, whereas MetRaC yields a distribution with confidence intervals. While straightforward to compute, constant rates do not capture the trajectory between measurements, lack uncertainty quantification, and are sensitive to measurement noise. Nevertheless, they offer a useful baseline for evaluating the advantages of the Bayesian approach. To enable a direct comparison with MetRaC, constant rate estimates were supplemented with bounds defined analogously to a confidence interval:

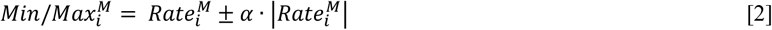

where α is a user-defined scaling factor. This formulation creates an interval around each constant rate estimate, allowing direct comparison with the Bayesian confidence intervals generated by MetRaC.

### 2.4 MetRaC-driven GEM reduction pipeline

The reduction pipeline begins with a generic model setup, followed by five sequential reduction steps: (1) detection and resolution of infeasibilities, (2) MILP-based selection of essential exchange reactions, (3) removal of redundant transport reactions, (4) elimination of reactions unused in parsimonious flux distributions (i.e., pFBA), and (5) removal of thermodynamically infeasible cycles. After each reduction step, dead-end metabolites and blocked reactions - arising from the removal of reactions - are systematically identified and pruned using Flux Variability Analysis (FVA) (Mahadevan and Schilling, 2003). This iterative procedure ensures that the reduced model remains feasible, minimal, and consistent with the imposed constraints.

#### 2.4.1 Model setup

The iCHO1766 GEM (Hefzi et al., 2016) was curated to match the cell line specificities. Auxotrophies specific to CHO-DG44, as reported in Hefzi et al. (2016), were enforced by closing the corresponding reactions (flux bounds set to zero). Specifically, the following enzymes were inactivated to reproduce amino acid auxotrophies:

- **Arginine**: argininosuccinate synthase (ARGSS),
- **Cysteine**: cystathionine gamma-lyase (CYSTGL) and cystathionine beta-synthase (CYSTS),
- **Proline**: glutamate-5-kinase (GLU5Km) and L-ornithine:2-oxo-acid aminotransferase (ORNTArm),
- **Lysine**: biotinidase (BTND1).

For all metabolites present in the culture media, both exchange and transport (between the extracellular and cytosol compartments) reactions were ensured to be present in the model. In the case of fumarate and fucose, which were present in the medium but absent from the extracellular compartment of iCHO1766, these reactions were explicitly added. Uptake was enabled only for metabolites present in the culture medium, ensuring realistic substrate availability.

Model constraints were introduced in two successive stages. A fixed set of exchange bounds was imposed once during model initialization. This included 58 medium metabolites: 28 core metabolites (acetate, alanine, arginine, asparagine, aspartate, cysteine, cystine, formate, fumarate, glucose, glutamine, glutamate, glycine, histidine, isoleucine, lactate, leucine, lysine, methionine, ammonium, phenylalanine, proline, serine, succinate, threonine, tryptophan, tyrosine, and valine), the recombinant product (measured daily in this study), and 29 additional components that were not measured experimentally in these cultures. Flux bounds for these metabolites were defined using the minimum and maximum uptake/secretion rates observed across multiple historical datasets (data not shown). In addition, several essential cofactors were always permitted for uptake and secretion, regardless of measurement availability. These comprised protons, water, phosphate, carbon dioxide, oxygen, sulphate, and a small set of ions (Fe^2+^, Na^+^, Ca^2+^, Cl^−^, HCO_3_^−^, and K^+^). A full list of constrained metabolites and their bounds is provided in Supplementary Table S1.

Second, for each measurement time point, the model was constrained with the calculated metabolic rates of the 28 core metabolites, together with the observed specific growth rate and recombinant product secretion rate. Importantly, these constraints were applied iteratively during the reduction process: at each reduction step, the strategy was evaluated across all available time points using the corresponding data. This ensured that experimental variability was consistently accounted for and that the reduced model remained feasible under the full set of dynamic conditions.

#### 2.4.2 Step 1 - Detect and resolve infeasibilities

At each experimental time point, the reduced model was subjected to a feasibility check under the applied constraints. If a valid solution existed, no adjustment was needed. If infeasibility arose, it indicated that the constraints associated with this experimental time point were overly restrictive. To diagnose and resolve such cases, we implemented a slack-variable MILP formulation, following a strategy similar to Klamt and von Kamp (2022) which identifies and relaxes conflicting constraints in a minimally invasive manner. Slack variables were added only to exchange reactions, permitting minimal adjustments to their bounds without altering the internal network structure. The MILP objective was to minimise the total slack introduced, ensuring that feasibility was restored with the smallest possible deviation from the experimental constraints:

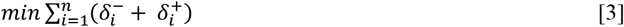

Subject to:

- Mass balance constraint: *S* · *v* = 0
- Lower bound constraint: 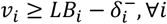
- Upper bound constraint: 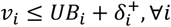
- Non-negativity of slack variables: 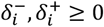
- Slack is only permitted on exchange reactions: 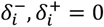, *if i* ∉ *exchange reactions*

where *i* indexes the reactions out of *n* total, *S* is the stoichiometric matrix. *v* is the flux vector, and *LB*_*i*_ and *UB*_*i*_ are the lower and upper bounds for each reaction *v*_*i*_. The slack variables, 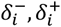 indicate the minimal adjustment required to restore feasibility.

Exchange reactions requiring non-zero slack were flagged as overly restrictive. To correct these cases consistently across the dataset, auxiliary demand reactions were introduced with bounds set to the minimal slack needed across all time points. This strategy ensured global feasibility while preventing excessive relaxation of individual constraints. All optimisation problems were solved using the open-source *HiGHS* solver (Huangfu and Hall, 2018).

#### 2.4.3 Step 2 - MILP-based exchange reactions selection

The iCHO1766 model includes over 600 exchange reactions, many of which are unnecessary for maintaining feasibility under the experimental conditions. To identify the minimal subset, we implemented a MILP in which each candidate exchange reaction was associated with a binary decision variable, indicating whether the reaction was retained (1) or removed (0) (Burgard et al., 2001; Reed and Palsson, 2004). The MILP problem was defined as follows:

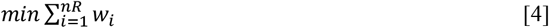

Subject to:

- Mass balance constraints: *S* · *v* = 0
- Flux upper bounds for removable reactions: *v*_*i*_ ≤ *w*_*i*_ · *UB*_*i*_ ∀*i* ∈ *removable*
- Flux lower bounds for removable reactions: *v*_*i*_ ≥ *w*_*i*_ · *LB*_*i*_ + *ε* ∀*i* ∈ *removable*
- Elimination of previously found solutions (to identify alternative minimal sets): 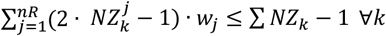

Here, *i* indexes the exchange reactions in the *removable* set, which includes all exchange reactions except the ones already constrained during the model setup (section 2.4.1). *v*_*i*_ denotes the flux through reaction *i*, and *w*_*i*_ is a binary variable indicating if the reaction is active (*w*_*i*_ = 1) or inactive (*w*_*i*_ = 0). *S* is the stoichiometric matrix, and *UB*_*i*_ and *LB*_*i*_ are the upper and lower flux bounds derived from experimental data. A small positive constant *ε* is added to the lower bound constraint to avoid numerical issues with strict zeros. *NZ* represents a previously identified solution and is used to exclude duplicate solutions during iteration. The MILP was solved separately for each time point, and the union of all exchanges identified as active was retained. Extracellular metabolites not linked to any retained exchange were removed along with their associated reactions.

#### 2.4.4 Step 3 - Removal of redundant transport reactions

Extracellular metabolites in GEMs are frequently linked to multiple possible transport mechanisms (e.g., passive diffusion, proton- or sodium-coupled transport). We curated a minimal set of transport reactions, applying a preference hierarchy: passive transport was retained whenever possible, followed by proton-coupled transport, and finally sodium-coupled transport. Alternative or duplicate transport routes were eliminated.

#### 2.4.5 Step 4 - Removal of reactions not used in parsimonious flux distributions

Parsimonious flux balance analysis (pFBA) (Lewis et al., 2010) was applied across all experimental time points to identify minimal-flux solutions. Reactions that did not carry flux in any of the parsimonious flux distributions were removed.

#### 2.4.6 Step 5 - Removal of thermodynamically infeasible cycles

As a final reduction step, we eliminated potential internal cycles permitting flux without net substrate consumption. To this end, we applied a loopless FBA filtering strategy (Desouki et al., 2015). At each time point, the model was constrained with the time-specific uptake and secretion bounds and then optimised for biomass and product (IgG) formation using both standard and loopless FBA. Reactions carrying flux only in the standard formulation were flagged as potential cycle participants. Reactions consistently identified as cycle participants under both objectives and across all time points were eliminated. This approach ensured that the reduced model remained free of infeasible flux loops while retaining essential functionality.

### 2.5 Transcriptomics data

In-house RNA-seq data of CHO-DG44 provided transcript per million (TPM) values for 16,383 genes across 192 samples, including 1,089 metabolic genes part of the iCHO1766 model. Expression values were transformed as log2(TPM + 1) to stabilise variance (VST). Essential genes were determined experimentally for the same cell line and are available in a public repository (Marzluf, 2025). Of 2,794 genes in this dataset, 308 were successfully mapped to NCBI identifiers, of which 233 corresponded to metabolic genes represented in the iCHO1766 model.

Transcriptomics-driven reduction was performed using the GIMME algorithm (Becker and Palsson, 2008). GIMME constructs a single context-specific model by retaining reactions required to achieve near-optimal growth while penalising reactions below the expression threshold. In our implementation, flux bounds were constrained to the overall minimum and maximum uptake/secretion values measured across all time points in the experimental dataset (model setup constraints – section 2.4.1). Our implementation of the algorithm integrates the transcriptomics data as a whole, rather than on a time-series basis, resulting in a reduced model for the dataset.

A common practice in transcriptomic analyses is to apply a fixed, data-independent cutoff, for example considering 1 VST unit (≈1 TPM on the log2(TPM+1) scale) as a threshold for weak expression. However, such global cutoffs risk misclassifying gene activity, as they do not reflect the specific distribution of expression values in a given dataset. To derive a more data-driven threshold, we examined the distribution of non-zero expression values (Richelle et al., 2019b). The 10^th^ percentile (0.05 VST) was too close to background noise to be reliable, so we adopted the 25^th^ percentile (4.03 VST) as our global cutoff for distinguishing expressed from weakly or non-expressed genes. Gene– reaction mapping was performed using the iCHO1766 gene–protein–reaction (GPR) rules, applying Boolean logic such that AND relationships were assigned the minimum expression across subunits, and OR relationships the sum across isozymes.

### 2.6 Metabolic task calculation

Metabolic functionality was assessed using the curated metabolic task framework (Richelle et al., 2021, 2019a). Each task specifies a minimal input–output metabolic conversion, such as the ability to synthesise an amino acid from defined precursors. For each evaluation, all exchanges were initially closed, and only the reactions associated with the substrates and products required for that task were allowed to carry flux. If a metabolite needed for the task was not already associated with an exchange reaction in the model, a temporary exchange reaction was added to allow its uptake or secretion for the task evaluation. To avoid spurious infeasibilities, a small set of cofactor exchanges was also left open (i.e., H^+^, H_2_O, Pi, Na^+^, PPi). A task was considered passed if the model resulted in a feasible solution. Tasks designed to be infeasible, for example, synthesizing essential amino acids for growth, were counted as passed when they failed. Out of 210 tasks, we evaluated 155; the others were excluded because their associated metabolites were not detected in our dataset (see Supplementary Table S2).

## 3. Results

### 3.1 Reduction of iCHO1766 using Bayesian MetRaC-derived metabolic rates

We applied the reduction pipeline to the CHO genome-scale model iCHO1766 (Hefzi et al., 2016), using time-course exo-metabolomic data from 12 fed-batch cultures. Metabolic rates were estimated with the Bayesian MetRaC framework (Andersson et al., 2025; Richelle et al., 2025), which provides uptake and secretion profiles with quantified confidence intervals (CIs). We first evaluated how different CI thresholds affect network structure, feasibility, and functional fidelity, and then applied the stepwise pipeline to generate a compact reduced model tailored to CHO bioprocess conditions.

To assess the role of CI thresholds on the reduced network structure, we generated four iCHO1766 variants by constraining exchange bounds with MetRaC rates at 68%, 95%, 99%, and 100% confidence intervals (CIs). The most substantial reduction occurred at 95% CI (Figure 1A), whereas wider bounds (99–100%) forced the reintroduction of reactions to maintain feasibility, preventing further reduction. This reflects a balance: moderately strict bounds remove flux routes unsupported by data, while overly wide bounds allow outliers to reopen exchange directions that had been closed under tighter constraints, increasing model size without improving its prediction capability.

**Figure 1.**
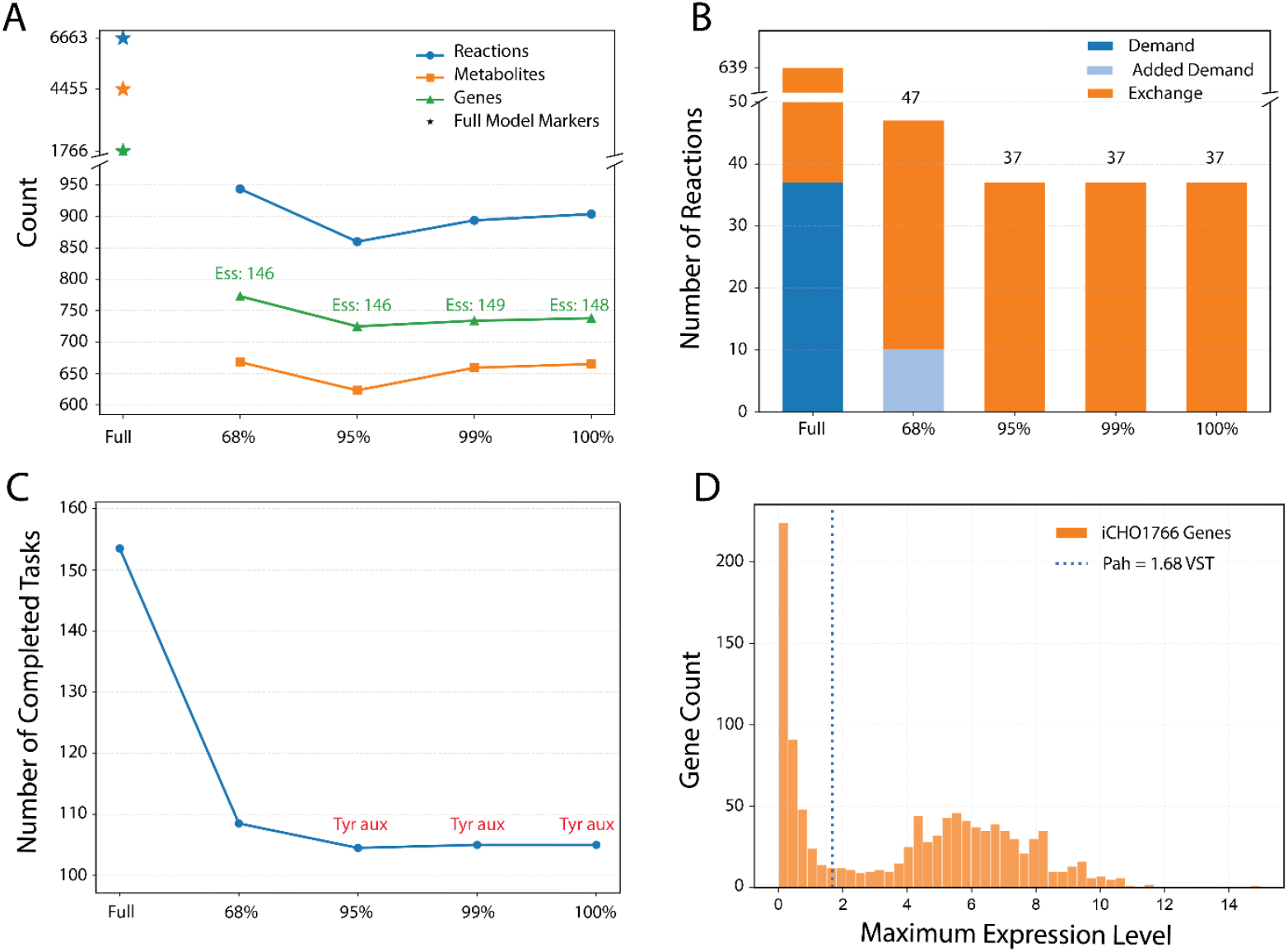
Impact of MetRaC confidence interval thresholds on iCHO1766 reduction. (A) Network size decreases most substantially at 95% CI, while stricter thresholds (99–100%) require reintroduction of reactions to maintain feasibility. (B) All reduced models converge to a minimal set of 37 exchange reactions; only the 68% CI model required ten additional demand reactions. (C) The number of completed metabolic tasks drops sharply at 68% CI and stabilizes around 105 of 155 tasks for 95–100% CI. Tyrosine auxotrophy consistently emerges from 95% CI onward. (D) Distribution of maximum gene expression values (Variance Stabilizing Expression, VST) in iCHO1766. Loss of tyrosine biosynthesis capacity coincides with low *PAH* expression (1.68 VST)

Exchange and demand reactions followed the same trend: all reductions required 37 essential exchanges, while only the 68% CI model required ten additional demand reactions to ensure system feasibility under the imposed experimental constraints (Fig. 1B). Metabolic task evaluation (as described in the Material and Methods) was used to confirm that core metabolic functions were generally preserved (Fig. 1C), although tyrosine synthesis from phenylalanine consistently failed from 95% CI onward. Low expression of *PAH* (1.68 VST) provides a plausible explanation for the observed auxotrophy, as the reaction could not sustain flux (Fig. 1D). Overall, the 95% CI provided the best compromise, yielding the smallest feasible network while retaining most metabolic functions.

Building on this observation, the full reduction pipeline was applied using the 95% CI-constrained model. Table 1 summarizes how structural size (reactions, metabolites, and genes) and functional performance (essential genes and metabolic tasks) evolved across successive reduction steps (i.e., resolving infeasibilities, MILP exchange reactions pruning, transport reactions clean-up, pFBA reduction, and loop removal). The initial model setup and infeasibility resolution (Step 1) reduced the network size by more than one-third while maintaining 136 of 155 metabolic tasks. There were no data-caused infeasibilities when using the 95% CI of the MetRaC data. Essential genes lost at this stage were associated with reactions blocked under the imposed constraints. Exchange reactions pruning (Step 2) eliminated unconstrained exchanges not needed to maintain model solvability at any experimental data points, converging to a minimal set of 37 exchange reactions. This set corresponded precisely to the 28 measured core metabolites, the recombinant product and growth rate constraints, plus a small group of essential exchanges (CO_2_, H, H_2_O, HCO_3_, O_2_, SO_4_, and Pi). The task feasibility decreased to 131/155.

**Table 1.** Structural and functional properties of the iCHO1766 model across successive reduction steps using 95% CI MetRaC rates.

| Model Reduction Steps | Reactions | Metabolites | Exchange Reactions | Genes | Essential Genes | Metabolic Tasks |
| --- | --- | --- | --- | --- | --- | --- |
| **Initial Model** | 6663 | 4455 | 602 | 1766 | 233 | 153/155 |
| **Step 1: Resolve Infeasibilities** | 4320 | 2228 | 282 | 1488 | 211 | 136/155 |
| **Step 2: MILP Exchanges** | 3566 | 1798 | 37 | 1366 | 195 | 131/155 |
| **Step 3: Transport** | 2174 | 1411 | 37 | 1139 | 185 | 130/155 |
| **Step 4: pFBA** | 860 | 623 | 37 | 725 | 146 | 105/155 |
| **Step 5: Thermodynamic Infeasible Loops** | 860 | 623 | 37 | 725 | 146 | 105/155 |

Transport reactions cleanup (Step 3) further reduced redundancy while pFBA-based trimming (Step 4) reduced the network by retaining only reactions that contributed to feasible parsimonious solutions, with inactive reactions systematically removed. This yielded a compact model of 860 reactions, 623 metabolites, and 725 genes, with 105 tasks retained over 155. Finally, thermodynamic infeasible loop removal (Step 5) left the structure unchanged, confirming that the prior reductions had already eliminated loop artifacts. Thus, the pipeline produced a condition-specific reduced network that retained all known auxotrophies and essential gene sets while drastically reducing model complexity. Note that tyrosine became essential because the phenylalanine-to-tyrosine (PAH) pathway became inactive under the data constraints (supported by transcriptomic data evidence, Fig. 1D).

The final model retained essential predictive capabilities while achieving substantial compression. Nearly nine out of ten reactions in the original model were eliminated (87% decrease), with the largest decreases in exchange/transport (−92%), lipid metabolism (−90%), and amino acid metabolism (−73%), while central subsystems for sustaining growth such as nucleotide metabolism (−62%) and energy metabolism (−34%) were more preserved (Fig. 2A). Importantly, the productivity–growth trade-off prediction was identical between the original and reduced models (Fig. 2B), demonstrating that compression did not compromise the prediction of key phenotypes.

**Figure 2.**
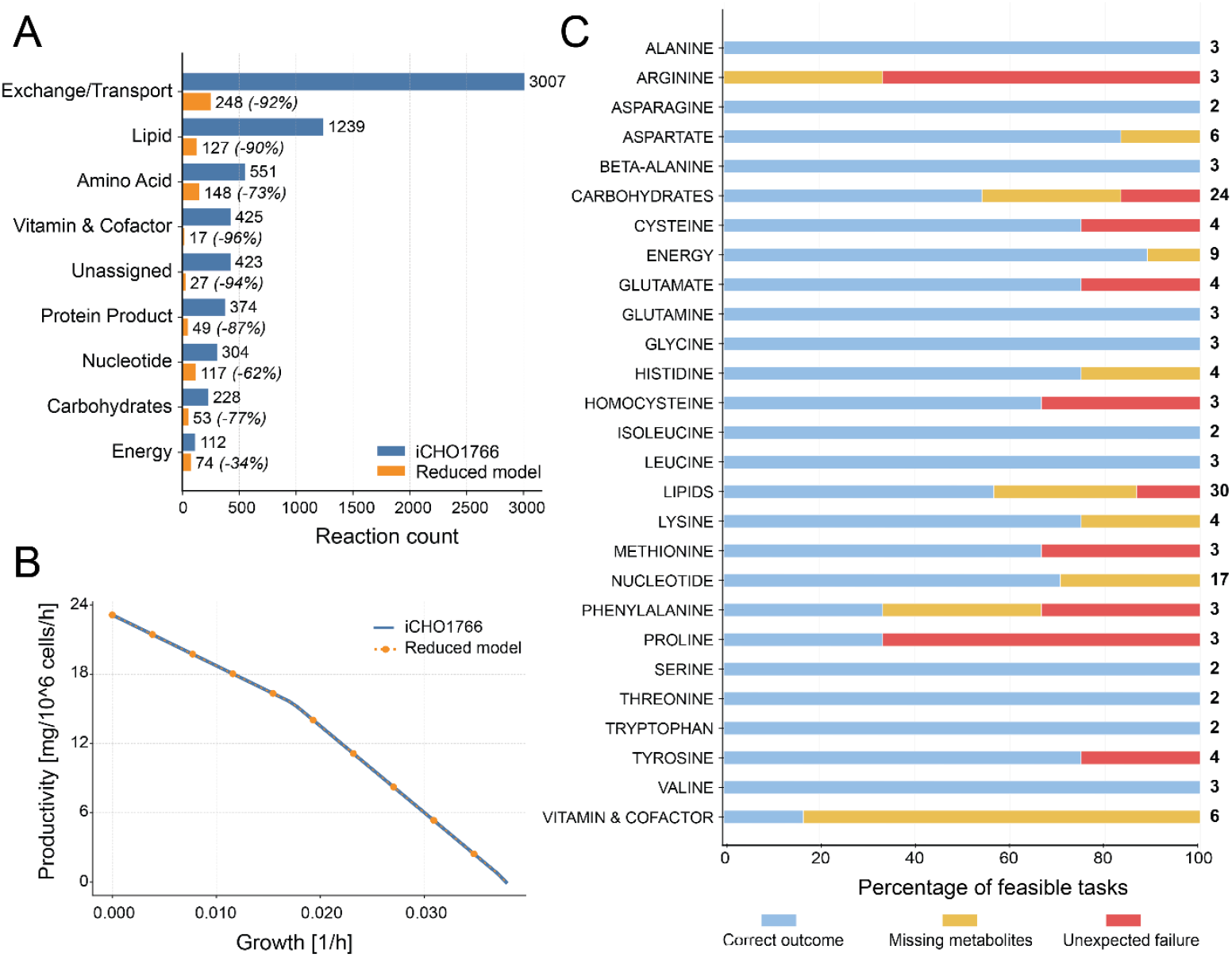
Effects of model reduction on network structure, phenotype, and task feasibility. (A) Reaction counts across subsystems in the original iCHO1766 model and the reduced version (95% CI). Substantial reductions were observed in exchange/transport, lipid, and carbohydrate metabolism, while protein product synthesis remained largely preserved. (B) Productivity–growth trade-off curves were unchanged, indicating that the reduced model maintained the same global performance envelope as the full model. (C) Outcomes of metabolic task testing for the 155 evaluated tasks. Each bar shows the fraction of feasible tasks per metabolite. The black numbers on the right side indicate the total number of tasks under each metabolic category. Blue: correct outcome (tasks that were feasible or failed by design); yellow: infeasible due to missing metabolites after reduction; red: unexpected failure (tasks expected to succeed but lost after reduction).

Metabolic task testing provided a detailed view of functional coverage (Fig. 2C). Out of 155 evaluated tasks (linked to metabolites detected in our dataset; Supplementary Table S2), 32 failed because the corresponding metabolites were absent from the reduced model. The remaining 18 were classified as “unexpected failures”. Of these, 8 were attributable to enforced DG44 auxotrophies for arginine (tasks 65, 67), proline (tasks 90, 128, 130), and cysteine (tasks 83, 105, 116). The other 10 failures stemmed from the reduction process itself: one after the removal of blocked reactions (task 175) and nine during the pFBA-based pruning step, where secondary pathways not required in parsimonious flux distributions were eliminated (tasks 27, 28, 30, 52, 124, 142, 165, 166, 167).

### 3.2 Impact of rate estimation method on reduced network features

We next compared the reduction pipeline using flux constraints from two estimation strategies: our uncertainty-aware MetRaC framework and a standard constant rate calculation method (see Materials and Methods). This comparison highlights how rate estimation shapes the reduced network. Unlike MetRaC, which explicitly integrates data noise and scarcity into probabilistic bounds, the constant rate method ignores uncertainty and relies on fixed point estimates. To approximate the role of confidence intervals, we introduced an α parameter that defines upper and lower bounds around each constant rate estimate.

Constraining iCHO1766 with constant rate estimates consistently produced larger reduced models than with MetRaC-derived rates, independent of the α value (Fig. 3A). Unlike MetRaC, where exchange requirements scaled predictably with confidence intervals, constant rate reductions showed no clear trend with α (Fig. 3B). We therefore focused on α = 0.3, the most constrained case, which still yielded a substantially larger network than the MetRaC reduction, with excess reactions in transport, nucleotide, and lipid metabolism (Fig. 3C).

**Figure 3.**
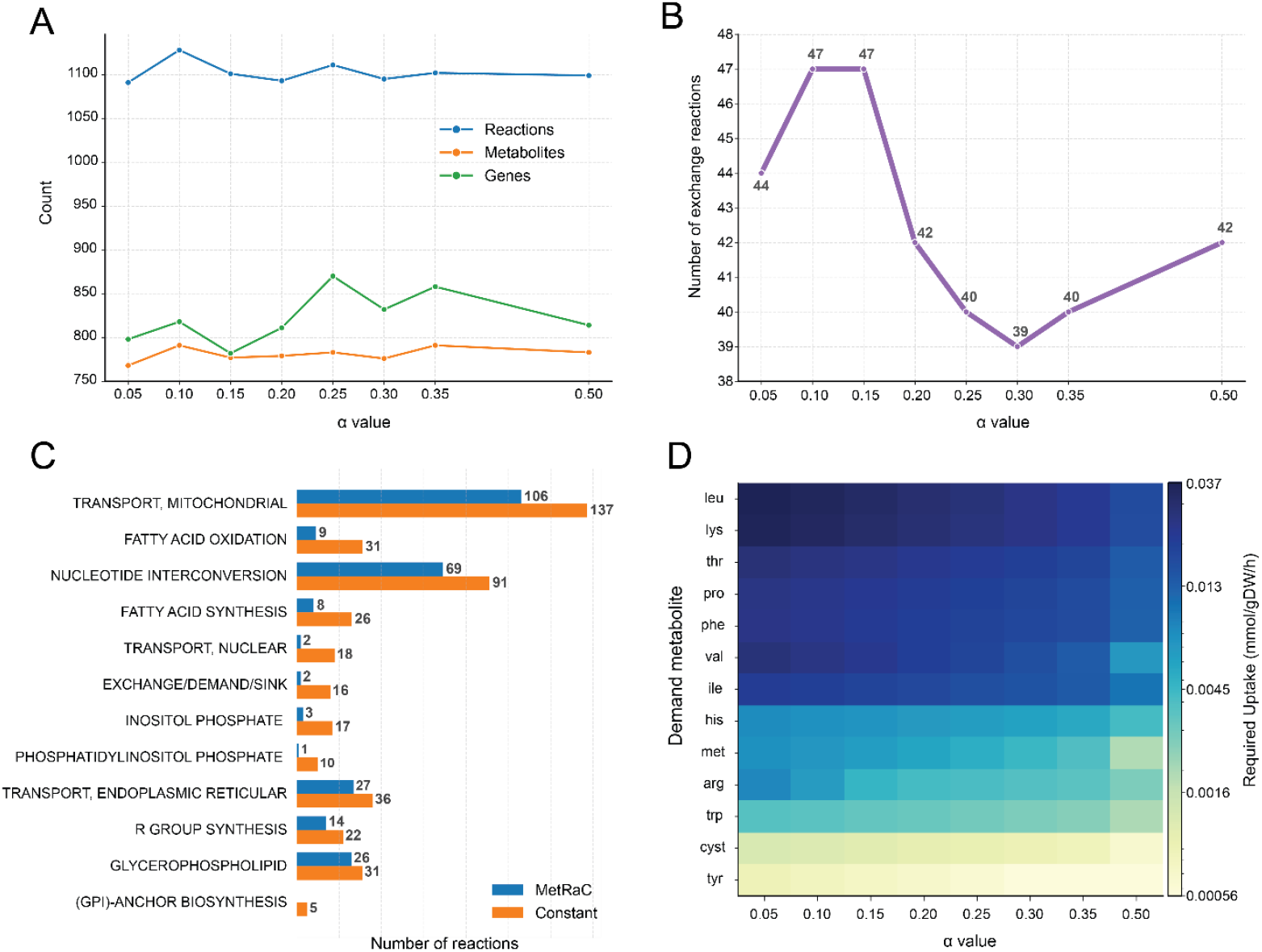
Model reduction using constant rates leads to increased model size and demand requirements to ensure feasibility. (A) Structural model size (reactions, metabolites, and genes) as a function of α. Constant rate reduction consistently produced larger networks than the MetRaC-reduced counterpart. (B) The number of exchange reactions fluctuated unpredictably with α. (C) Subsystem distribution of retained reactions at α = 0.3 highlights the increased number of reactions in constant rate reductions relative to the MetRaC-reduced model. (D) All constant rate reductions required demand reactions to ensure feasibility across all time points. Although uptake magnitudes decreased with increasing α, several amino acids still required additional uptake, highlighting the limitations of constant rate estimation.

Feasibility was another major limitation. Across all α values, constant rate reductions required additional demand reactions to restore feasibility (Fig. 3D). Although the magnitude of these artificial uptakes declined as α increased, several amino acids remained dependent on non-biological demand fluxes. This reflects a key inconsistency: constant rate models were forced to assume metabolite uptake not supported by the experimental data. By contrast, MetRaC approach maintained feasibility without additional metabolic constraints.

Direct comparison of rate profiles (Supplementary Fig. S1) revealed the most likely source of these issues: constant rates frequently produced spurious spikes and biologically implausible secretion of essential amino acids. Together, these results show that constant rate reductions yield larger and less stable networks, while MetRaC reductions remain smaller, feasible without artificial adjustments, and biologically consistent thanks to the availability of time-resolved, uncertainty-aware bounds.

### 3.3 Metabolomics-guided reduction retains transcriptomics-consistent functionality

To evaluate whether metabolomics-only reduction can preserve functions typically captured by gene-expression–guided strategies, we compared our MetRaC-based model (95% CI ) with a transcriptomics-driven reduction obtained using GIMME (Becker and Palsson, 2008). GIMME primarily retains reactions supported by highly expressed genes, under the assumption that transcriptional activity reflects metabolic relevance, while still relying on generic exchange constraints. In contrast, our metabolomics-guided reduction is based solely on feasibility under time-resolved extracellular flux data, without any transcriptomic input. This comparison allowed us to test whether metabolomics constraints alone are sufficient to recover intracellular functions that align with gene expression evidence.

The two reduced models shared 488 metabolites, 551 reactions, and 509 genes (Fig. 4A), but diverged in both structure and function. The GIMME model was substantially larger yet completed fewer metabolic tasks than the MetRaC model (Table S3). This counterintuitive result stems from differences in how constraints act: transcriptomics-based filtering inflates network size by leaving many reactions unconstrained (due to missing gene–protein–reaction links) while discarding reactions linked to weakly expressed but functionally essential genes. Consequently, several pathways required for amino acid and central carbon metabolism failed in the GIMME model. In contrast, the metabolomics-driven reduction preserved feasibility across all experimental time points and maintained a higher fraction of metabolic tasks, despite relying only on extracellular data.

**Figure 4.**
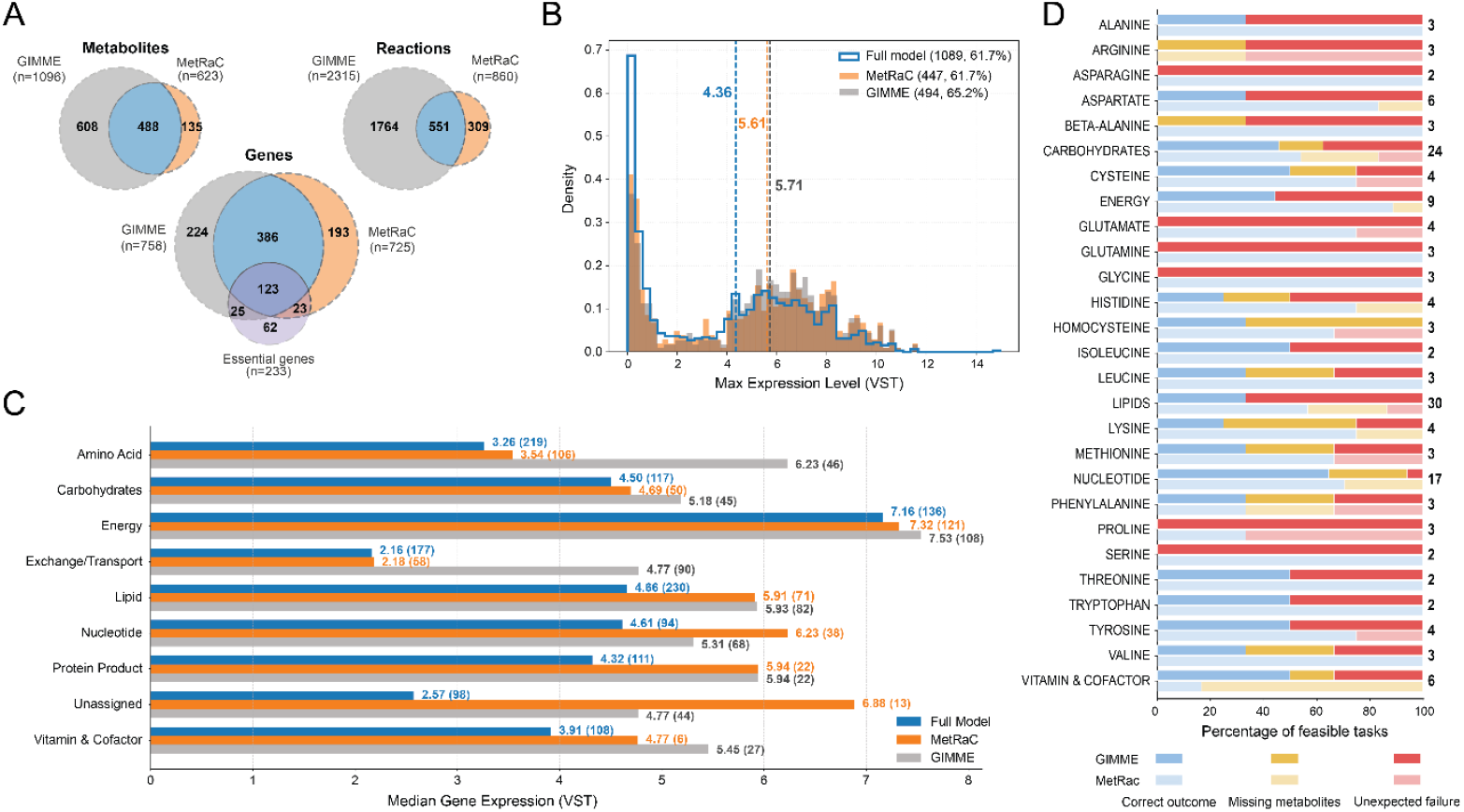
Structural and function differences in metabolomics- and transcriptomics-guided reductions. (A) Overlap of metabolites, reactions, and genes between transcriptomics-reduced (GIMME) and metabolomics-reduced (MetRaC, 95% CI) models. (B) Distribution of maximum gene expression levels (VST) in iCHO1766 and the reduced models. Both reductions removed low-expression genes, raising median expression (4.36 in iCHO1766 vs 5.61–5.71 in reduced models). (C) Median gene expression per metabolic subsystem and number of associated genes (number in brackets). (D) MetRac and GiMME outcomes of metabolic task testing for the 155 evaluated tasks. Each bar shows the fraction of feasible tasks per metabolite. The black numbers on the right side indicate the total number of tasks under each metabolic category. Blue: correct outcome (tasks that were feasible or failed by design); yellow: infeasible due to missing metabolites after reduction; red: unexpected failure (tasks expected to succeed but lost after reduction).

Both models retained a similar set of essential genes, consistent with their strong expression and central role in biomass production. Expression-level distributions confirmed that both approaches preferentially kept highly expressed genes, raising median expression compared to the full iCHO1766 model (4.36 vs. 5.61–5.71; Fig. 4B). The nearly identical medians highlight that the metabolomics-based reduction, despite not using transcript thresholds, preserved a comparable set of strongly expressed genes to GIMME. However, subsystem analysis revealed a key distinction (Fig. 4C–D): the metabolomics-guided reduction maintained broader functional coverage, whereas the transcriptomics model - despite retaining many highly expressed genes - failed numerous tasks in amino acid and central carbon metabolism (e.g., glucose, lactate, succinate).

Together, these results show that metabolomics-driven reduction, even without explicit transcriptomic input, can yield a compact yet functionally robust model that preserves concordance with gene expression patterns. This highlights its potential as a scalable alternative to transcriptomics-driven reductions, especially when intracellular data are unavailable.

## 4 Discussion

This study establishes a generalizable framework for deriving compact, uncertainty-aware metabolic models that can serve as plug-and-play components in digital twins for CHO bioprocesses and, more broadly, biopharmaceutical applications. The central contribution lies not only in the reduction pipeline itself but in how experimental evidence is integrated into the process. By leveraging the MetRaC framework for Bayesian inference of metabolic rates, we incorporate experimental uncertainty directly into constraint definition. This goes beyond traditional constant-rate approaches, which treat extracellular fluxes as static averages and often ignore method-specific error or the temporal variability of cellular metabolism. Incorporating time-resolved confidence intervals allowed us to derive reduced models that remained feasible across all experimental conditions without requiring artificial demand reactions and that were ∼20% smaller than constant-rate reductions.

The pipeline builds on these uncertainty-aware flux estimates through a structured sequence of reduction steps that progressively remove redundancy while preserving functionality. Diagnostics using slack-variable MILPs ensure that infeasible constraints are curated rather than blindly enforced, and non-necessary exchange, redundant transport, and zero-flux reactions are pruned systematically. Importantly, quality control is embedded throughout the process: after each pruning step, experimental constraints are reapplied, and feasibility is verified across all time points. This iterative safeguard prevents over-reduction and ensures that the resulting model reliably captures the observed dynamics. The outcome is an eight-fold compression of iCHO1766, from 6,663 to about 800 reactions, without compromising core phenotypic predictions such as the productivity-growth trade-off.

For digital twin applications, feasibility and computational efficiency are more critical than maximal biological completeness. A reduced model must reproduce observed phenotypes consistently under relevant process conditions, since infeasibility in even a single time point would compromise predictive use. Our metabolomics-driven framework addresses this practical requirement while still preserving a strong mechanistic basis. Compared with transcriptomics-guided reductions such as GIMME, which emphasize intracellular gene activity, our approach relies solely on extracellular fluxes. Surprisingly, despite not using gene expression information, the metabolomics-guided reductions retained a similar set of strongly expressed genes and achieved broader metabolic task coverage. This suggests that essential cellular functions can be effectively captured by phenotypic constraints alone. For industrial biotechnology, where transcriptomics data may be unavailable or impractical to generate for proprietary cell lines and processes, this finding highlights a pragmatic route toward building functional, predictive models.

Despite its strengths, the framework has several important limitations that warrant discussion. First, the approach is inherently dependent on the quality of exo-metabolomics data: systematic measurement biases or missing values for key metabolites can propagate through the rate estimates and affect downstream pruning, even with the feasibility safeguards in place. Any improvement in rate estimation - for example, by incorporating biologically informed priors, data reconciliation, or better measurement error models - can be plugged into the pipeline and would likely reduce such propagation effects.

Second, constraining the model only with extracellular fluxes limits our ability to capture intracellular regulatory phenomena that do not manifest at the exchange level. Integrating secondary omics (transcriptomics, proteomics) as complementary filters could mitigate this gap, but it also introduces trade-offs: additional data layers may correct some omissions while introducing new biases from their own measurement noise or annotation gaps. Systematic studies are needed to define a principled trade-off between what should be enforced by exo-metabolomics and what should be constrained by intracellular omics (e.g., through weighted constraints, consensus rules, or hierarchical integration strategies).

Third, in Step 1 (infeasibility resolution), feasibility was restored by introducing demand reactions whenever experimental constraints proved overly restrictive. While this pragmatic fix enables the reduction to proceed, it inevitably expands the solution space and may allow fluxes not biologically supported by the data. Future work could explore more targeted alternatives, such as selective correction or reconciliation of problematic measurements, or softening infeasible bounds via penalised slack terms rather than introducing artificial uptake. These strategies would help preserve feasibility while reducing bias in the resulting reduced networks.

Finally, the pipeline currently applies each reduction step iteratively across all time points, aggregating results into a final solution. While this ensures that feasibility is respected under all experimental conditions, it can create unnecessary computational cost and lead to less parsimonious outcomes. For example, in Step 2 (MILP-based exchange reaction selection), the iterative strategy identifies the minimal set of exchanges required at each individual time point and then takes the union of all these sets. This conservative approach guarantees feasibility but often retains more exchange reactions than strictly necessary across the experiment. More efficient alternatives could involve parallelising the time-point MILPs or formulating a single optimisation problem that directly seeks the smallest set of exchanges ensuring feasibility across all time points simultaneously. Although illustrated here for Step 2, similar considerations apply to other reduction stages where per-time-point aggregation is used.

Together, these limitations point to opportunities for refinement - improving rate estimation, integrating omics layers more judiciously, streamlining the iterative steps, and replacing demand-reaction fixes with biologically grounded alternatives. Addressing these challenges will enhance the robustness and generalizability of the pipeline, particularly for large-scale or real-time applications.

In summary, we demonstrate that metabolomics-guided reduction produces compact, uncertainty-aware metabolic models that retain biological relevance and are well-suited for digital twin applications. The resulting models are computationally efficient for real-time simulation yet maintain the mechanistic underpinnings necessary for process optimization and control. By bridging experimental metabolomics with systematic model reduction, this framework moves genome-scale metabolic modeling closer to practical deployment in bioprocess digitalization.

## Acknowledgements

We thank Anton Vernersson, David Andersson, and Shanti Pijeaud for their help with the rate calculations, and the Sartorius Corporate Research team for their support in developing this study.

