## Supplementary figures and images for "Systematic data-driven genome-scale metabolic model reduction for dynamic bioprocess modeling: CHO cell culture case study"

### Supplementary Figure

**Figure S1** - Rate Comparison – MetRaC Rates 95% CI vs Constant Rates

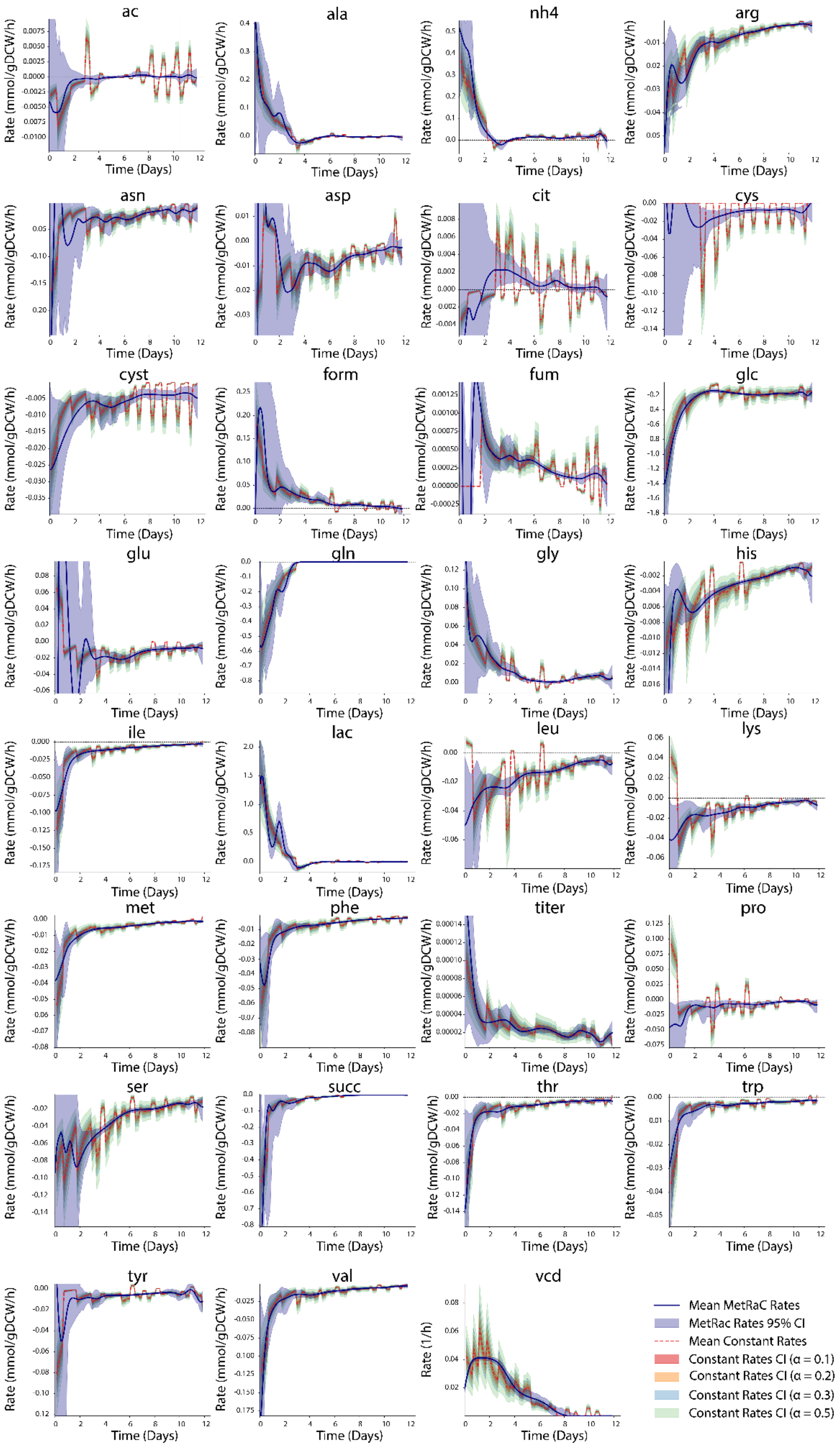
